# Ultrastructure of immature synaptic inputs in the lateral superior olive of rodent brainstem

**DOI:** 10.1101/2024.08.09.606139

**Authors:** Siyi Ma, Shigeki Watanabe, Deda C. Gillespie

## Abstract

Neurons of the lateral superior olive (LSO), which compute intensity differences between the two ears, receive two primary inputs, an ipsilaterally arising excitatory input and a contralaterally arising inhibitory input, that are precisely matched for stimulus frequency. Circuit refinement to establish this precise match takes place within the first few postnatal weeks through elimination of single-fiber inputs and concomitant strengthening of the remaining inputs. However, little is known about the ultrastructure of these young synapses and about how changes in physical features of these synapses could contribute to refinement. To characterize pre-hearing postnatal development of somatic synapses in the LSO, we performed transmission electron microscopy and examined synapses in the rodent LSO from birth to hearing onset at postnatal day 13. Synaptic vesicles and mitochondria in putative synaptic boutons were surprisingly scarce at birth. During the second week, bouton enlargement was accompanied by an increase in the number of vesicles and mitochondria. The size of mitochondria also increased, pointing to changes in functional and metabolic needs of synapses. Our results reveal extensive remodeling at individual presynaptic terminals that could strengthen single-fiber inputs and contribute to the development of robust synaptic transmission.

## Introduction

The lateral superior olive (LSO) of auditory brainstem receives primary excitatory and inhibitory inputs that arise from the ipsilateral and contralateral ears. The excitatory and inhibitory inputs form topographic maps, and these two maps must arrive at the LSO precisely registered. Indeed, the computation of interaural level differences by individual LSO principal neurons requires that their inputs be precisely matched for stimulus frequency (Tollin, 2003). In rats, the primary glutamatergic input from the ventral cochlear nucleus (VCN) and the primary glycinergic input from the medial nucleus of the trapezoid body (MNTB) innervate the LSO as early as embryonic day 18 (E18) (Kandler and Friauf, 1995). During the first two postnatal weeks, prior to hearing onset at about postnatal day 12 (P12), both MNTB and VCN pathways undergo physiologically measurable refinement that is characterized by the elimination of axonal fibers and the strengthening of the remaining inputs (Case et al., 2011; Kim and Kandler, 2003; Kim and Kandler, 2010). As has been demonstrated in the inhibitory projection, pre-hearing refinement depends on activity spontaneously generated in the immature cochlea (Clause et al., 2014; Tritsch et al., 2007).

Functional refinement has been more extensively studied in the MNTB-LSO pathway than in the VCN-LSO pathway. In mice, functional strengthening has been estimated to comprise a 2-fold increase in quantal size and an 8-fold increase in quantal content (Kim and Kandler, 2010) and is mostly completed before hearing onset (Walcher et al., 2011). Functional refinement is accompanied by a shift in the relative proportion of GABAergic and glycinergic components – from primarily GABAergic to primarily glycinergic transmission – due to changes in neurotransmitter release and postsynaptic receptor expression in the first two postnatal weeks (Korada and Schwartz, 1999; Kotak et al., 1998; Nabekura et al., 2004). Structurally, the addition of boutons to individual MNTB fibers during this period suggests that an increase in synapse number contributes to functional strengthening (Clause et al., 2014). Although structural changes at the level of a single synapse can also contribute to functional strengthening via increases in quantal content (Kaeser and Regehr, 2017; Vos et al., 2010), ultrastructural studies in the LSO to date have focused on juvenile and adult animals (Gjoni et al., 2018; Helfert et al., 1992), and the extent to which ultrastructural plasticity or maturation could underlie functional strengthening prior to hearing onset has not been examined.

Immature inhibitory synapses in the LSO are of especial interest because before hearing onset, MNTB terminals transiently express the vesicular glutamate transporter 3 (VGLUT3), enabling the release of not only GABA and glycine, but also glutamate, onto principal neurons of the LSO (Gillespie et al., 2005). As individual MNTB cells release glutamate and GABA/glycine (Alamilla and Gillespie, 2011), and the relevant vesicular transporters for glutamate and GABA/glycine colocalize at the level of conventional light microscopy before hearing onset (Gillespie et al, 2005), a significant subset of immature inhibitory terminals must contain both glutamatergic and GABA/glycinergic vesicles. Whereas the critical role of VGLUT3 expression and glutamate release in normal developmental refinement of this inhibitory pathway (Noh et al., 2010) raises questions for development generally, the existence of transiently co-transmitting immature synapses also raises more general questions about synapse ultrastructure, especially in light of classical methods used to identify synaptic phenotype (Gray, 1969).

We acquired 2D electron micrographs centered on LSO principal neurons at three ages between birth and hearing onset (inclusive) and examined somatic synapses to look for ultrastructural correlates of functional refinement previously described in the LSO. Between P1 and P13, we found significant increases in synapse parameters associated with synaptic strength, including bouton and mitochondrion size, and vesicle and mitochondrion number. Furthermore, the population of somatic synapses shifted from being mostly excitatory to mostly inhibitory. In sum, axon rearrangement and developmental refinement in the LSO are accompanied by dramatic changes in the presynaptic terminal that could underlie increases in synapse strength and reliability.

## Methods

### Animals

All animal procedures were performed in accordance with the Canadian Council on Animal Care guidelines and were approved by the Animal Research Ethics Board of McMaster University. Sprague-Dawley rats ages P1 – P13, born on site, were used in this study without regard to sex. All rats were housed under 12/12 h light/dark cycles and tissue was collected between Zeitgeber Time (ZT) 2 and ZT 8.

### Electrophysiology

#### Tissue Preparation

Animals aged P2, P6, and P12 were anesthetized with isoflurane and decapitated. Brains were quickly extracted into oxygenated (using 95%O_2_/5%CO_2_) ice-cold dissection artificial cerebrospinal fluid (ACSF) containing (in mM) 124 NaCl, 5 KCl, 1.25 KH_2_PO_4_, 25 NaHCO_3_, 0.1 CaCl_2_, 3 MgCl_2_, and 1 kynurenic acid (KA). Coronal slices of the brainstem were cut at 250 μm on a vibrating tissue slicer (Compresstome VF-300, Precisionary Instruments). Slices containing the MNTB and LSO were collected and submerged in oxygenated recording ACSF for 30 minutes at 35°C, and thereafter at room temperature until recording. For recording, ACSF contained (in mM) 124 NaCl, 5 KCl, 1.25 KH_2_PO_4_, 25 NaHCO_3_, 2 CaCl_2_, 1.3 MgCl_2_, and 1 kynurenic acid.

#### Electrophysiological recordings

After recovery, slices were transferred to a recording chamber perfused with oxygenated recording ACSF at 32°C. Stimulating electrodes (1-2 MΩ glass pipettes) filled with ACSF were inserted into the fiber tract exiting the MNTB. Principal neurons of the target LSO were visualized with DIC-IR and identified by their morphology and orientation within the nucleus. Only cells in the medial and middle limbs of the LSO were targeted. Recording electrodes (2-4 MΩ) contained (in mM) 74 Cs-methanesulfonate, 56 CsCl, 10 Na_2_-phosphocreatine, 10 EGTA, 10 HEPES, 0.3 Na-GTP, 4 Mg-ATP, 0.1 spermine, 5 QX-314. Whole-cell patch clamp recordings of LSO neurons were acquired at 20 kHz, with 4 kHz filtering (Multiclamp 700, Digidata 1440, pClamp 10, Axon Instruments). Cells were held at −60 mV (uncorrected for calculated junction potential of ∼4 mV). A Master-8 (with IsoFlex SIU, AMPI) was used to deliver current pulses (10–100 μA, 0.3 ms) to the MNTB fiber tract every 20 seconds. Postsynaptic currents (PSCs) were considered for analysis only if they occurred within 2 ms of the MNTB stimulus. Recorded traces were analyzed using Clampfit and custom MATLAB scripts to find peak amplitude and to fit single-exponential decay time constants (tau).

### Electron Microscopy

#### Tissue Preparation

Animals aged P1/2, 6/7, 12/13 were anesthetized with isoflurane and decapitated. Brains were quickly extracted into oxygenated ice-cold dissection ACSF (as above). Coronal slices of the brainstem (150 μm) were cut at the Compresstome. Slices containing both MNTB and LSO were collected and submerged in an incubation chamber filled with oxygenated recording ACSF at 35°C. The slices were left to recover for at least 30 minutes at 35°C, then transferred for 10– 20 minutes to a humidified interface chamber at room temperature. Although we initially prepared living slices and filled LSO neurons with Lucifer Yellow for photo-conversion, we found that the LSO cell somata could be easily identified in TEM images without previous labeling. We therefore prepared tissue for EM without labeling first, so as to minimize tissue deterioration that might otherwise occur throughout the process of preparing and labeling living slices. Slices were fixed overnight in 2% paraformaldehyde/6% glutaraldehyde in 0.1 M sodium phosphate buffer (PB) at 4°C, and then kept in a mixture of 160 mL 0.05 M PB, 120 mL ethylene glycol and 120 mL glycerol (Bolam, 1992) until being processed for electron microscopy.

Coronal sections were bisected and then trimmed to create two sections of roughly 4 mm x 4 mm that each contained one LSO. The trimmed sections were thoroughly washed 3 times (10 minutes each) with 0.1 M cacodylate buffer (CB) and were postfixed with 1% osmium tetroxide and 1% potassium ferrocyanide in 0.1 M CB for 90 minutes on ice. Sections were then washed 3 times (10 minutes each) with ddH_2_O and incubated in 2% uranyl acetate (UA) in ddH_2_O for 60 minutes in the dark. Sections were next dehydrated with increasing concentrations of ethanol (30%, 50%, 70%, 90% x2, 100% x 3) for 10 minutes each on ice followed by infiltration with increasing concentrations of Epon-Araldite ethanol mixture (30%, 70%) for 2 hours each. For 70% Epon-Araldite incubation, sections were placed in polyethylene BEEM capsule caps. Sections were incubated in 90% Epon-Araldite overnight and placed in fresh 100% Epon-Araldite (Araldite 4.4 g; Epon 6.2 g; DDSA 12.2 g; and BDMA 0.8 ml) 3 times for 2 hours each before being left to cure at 60°C for 3 days. All steps preceding the 70% Epon-Araldite incubation were performed on a shaker. Once the resin had cured, the tissue was cut from the resin block and mounted on a cylindrical resin block using cyanoacrylate glue. Semi-thin sections (400 nm) were cut on the ultramicrotome using a glass knife and stained with toluidine blue. Once the LSO appeared in the toluidine blue-stained sections, the block was re-trimmed to contain solely the LSO. Thin sections (50 nm) were then collected using a Diatome diamond knife and mounted onto 0.7% pioloform-coated copper slotted grids.

#### Transmission electron microscopy (TEM)

Samples were imaged using a ThermoFisher Talos L120C equipped with a Ceta camera. TEM images of the LSO nucleus were captured at low magnifications (2000X to 4000X) and LSO principal cells were identified by their morphology and orientation within the nucleus. Once cells were identified, a higher magnification (pixel size = 825 pm) was used to capture synapses near the soma. Approximately 8 cells from the medial and middle limb were sampled from each section. Where possible, the same tissue block was sampled at a depth at least 15 μm distant to capture a novel cell. At least two biological samples were used for each age.

#### EM Image Analysis

Electron micrographs were analyzed using modified ImageJ/FIJI macros from SynapsEM and associated MATLAB scripts (Watanabe et al., 2020; https://github.com/shigekiwatanabe/lso_EM_manuscript). All somatic synapses from a total of 18 cells (6 cells per age) were analyzed for this study. Synapses from 2 cells per age were randomized into a single pool, and this process was repeated twice to created 3 randomized pools, each comprising 6 cells (2 from each age).

Putative synaptic boutons were identified by their shape, presence of vesicles, and proximity to the LSO soma (membrane contact < 20 nm gap). All putative boutons were accepted, and none were excluded based on size or synapse contact length. Due to the sparsity of vesicles at the youngest ages, synapse-like structures that contained no vesicles were also included in the analysis. The plasma membrane, active zone, synaptic vesicles, and mitochondria were annotated in ImageJ/FIJI. Boutons and mitochondria were annotated with a freehand selection tool to compute size (area). Each synaptic vesicle was first annotated with a freehand selection and then fitted with an elliptical shape. Parameters for all synaptic components were quantified using custom MATLAB scripts from SynapsEM. Two naïve observers were trained to annotate cells; their analyzed measurements were consistent with the trends shown in this paper.

Example micrographs shown here were adjusted for brightness and contrast for clarity using ImageJ/FIJI.

### Immunohistochemistry

#### Tissue Preparation

Six animals, two each at ages P1, P6, and P13, were deeply anesthetized with sodium pentobarbital (120mg/kg) and transcardially perfused with 0.1M phosphate buffered saline (PBS) followed by cold 4% paraformaldehyde in PBS. Brains were extracted, postfixed overnight in the same fixative, and cryoprotected in 30% sucrose in PBS until time of sectioning (≤ 2 months of perfusion).

#### Immunohistochemistry

Coronal sections containing the LSO were cut at 30 μm on a freezing microtome and collected into PBS. Sections were co-immunostained for synapsin, VGLUT1, and VIAAT. In some cases, a fluorescent Nissl stain (NeuroTrace 430/450) was included to visualize cells. Two staining runs – each run containing tissue from each of the 3 ages – were completed, for a total of 2 biological samples per age. All immunohistochemistry was performed on free-floating sections at 4°C. Tissue sections were incubated in blocking solution [5% normal donkey serum (NDS), 2.5% bovine serum albumin (BSA), and 0.5% Triton X-100 in PBS] for 15 hours, in primary antibodies [diluted in 5% NDS and 2.5% BSA in PBS] for 24 hours, in secondary antibodies [diluted in PBS] for 24 hours, and then in counterstain [NeuroTrace] for 30 minutes (Extended Data Table 7-1). Sections were thoroughly washed, 10 minutes each, between each incubation step and before mounting and coverslipping with ProLong Gold (Invitrogen, P36934). Slides were cured for at least 36 hours at room temperature in the dark before imaging. For each staining run, a condition omitting all primary antibodies and a condition omitting all secondary antibodies were included for experimental controls.

#### Image acquisition and analysis

Images were acquired with a confocal microscope (Zeiss LSM 980 AiryScan), using a 63X oil objective (NA = 1.4) to achieve a pixel size of 44 nm, with sequential imaging for each fluorophore. Analysis was performed using custom ImageJ/Fiji and MATLAB scripts. Principal neurons of the LSO were identified by their shape and orientation, and somata were selected manually; subsequent analysis of terminal staining around the soma was automated. First, each selection of the soma was dilated 1.5 μm to encompass presynaptic label around the soma. To segment synaptic label, each image channel was thresholded at a value 4 standard deviations (4 S.D.) above the mode of the pixel intensity distribution, a process that likely resulted in undersampling, but that was chosen to minimize artifactual overlap. A searching region with radius 200 nm from the centroid of each synapsin-identified synapse was then used to search for overlapping thresholded clusters of VGLUT or VIAAT label. Overlaps are reported as fraction of total synapsin^+^ clusters.

### Statistical Analysis

Scatterplots with error bars show medians with 25% and 75% quartiles. All graphs and statistical analyses were performed using GraphPad Prism 10. Comparisons among the three age groups were done using a rank-based nonparametric test (Kruskal-Wallis test) followed by nonparametric pairwise multiple comparisons (Dunn’s test). Exact p-values are reported to 4 significant digits except where p < 0.0001.

## Results

### MNTB inputs are eliminated, and remaining inputs strengthened, between P1 and P13

Inputs to the rodent LSO from both MNTB and VCN undergo refinement before hearing onset that is characterized by a decrease in the number of single-fiber inputs and an increase in the strength of remaining single-fiber inputs (Case et al., 2011; Kim and Kandler, 2003; Kim and Kandler, 2010). To correlate physiologically measurable refinement with ultrastructure at the specific ages used in this study, we made whole-cell recordings of postsynaptic currents (PSCs) from LSO neurons while gradually increasing electrical stimulation of the MNTB input pathway to recruit additional input fibers (Fig. 1). From the resultant input-output relationships, we estimated the strength of MNTB fiber inputs to individual LSO neurons at ages P1/2, P6/7, and P12/13. In replication of earlier work (Kim and Kandler, 2003), postsynaptic currents and input-output curves for individual P1/2 neurons (Fig. 1A) exhibited many small increases in PSC amplitude as a function of increasing stimulus intensity. Both P6 and P12 neurons showed fewer, but larger, discrete increments in response strength as stimulus intensity increased (Fig. 1B-C). Estimated minimum (medians = 0.07, 0.26, and 6.02 nA) and maximum response strength (medians = 0.58, 4.60, and 14.0 nA) both increased with age (n = 5 for P1/2, 4 for P6/7 and P12/13; Fig. 1D-E). The increasingly rapid decay kinetics (τ = 5.3 for P1/2; τ = 4.7 ms for P6/7; τ = 1.4 ms for P12/13) likely reflect an age-dependent shift from GABA-dominant to glycine-dominant transmission at these synapses (Korada and Schwartz, 1999; Kotak et al., 1998; Nabekura et al., 2004), whereas the appearance of discrete steps in the input-output curves of older slices, and the increases in estimated minimum and maximum PSC amplitudes, are consistent with the elimination of early inputs and the strengthening of remaining inputs (Kim and Kandler, 2003; Kim and Kandler, 2010).

**Figure 1.**
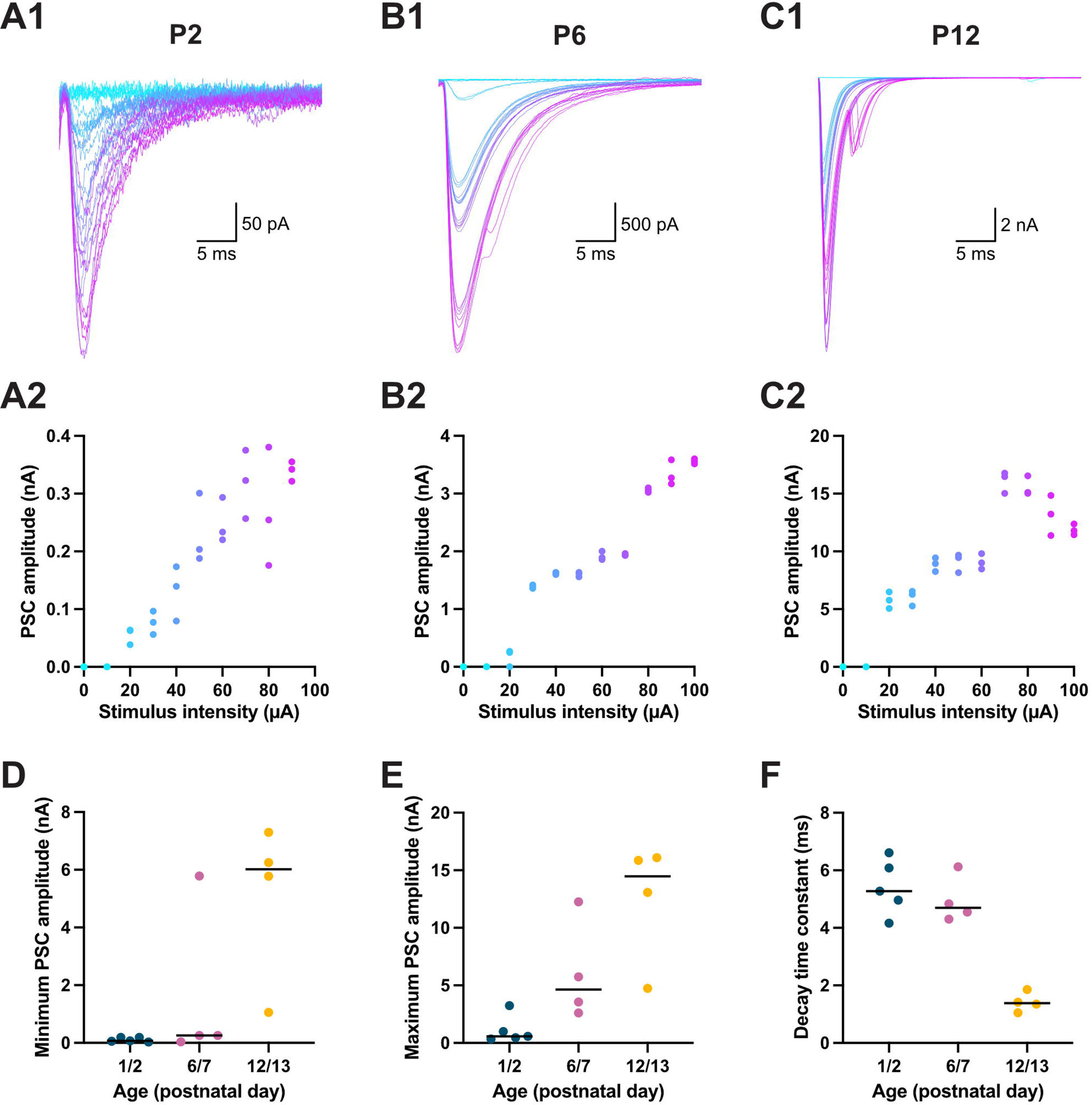
MNTB inputs to LSO grow in strength and decrease in number between P1 and P13. Representative PSCs (***A1, B1, C1***) and their corresponding input-output curves (***A2, B2, C2***) collected in individual LSO neurons in response to increasing electrical stimulation (cyan to magenta) in the MNTB pathway at P2, P6, and P12. ***D***) Estimated minimum fiber strength as a function of age (n = 5 for P1/2; 4 for P6/7 and P12/13; blue, pink, yellow). ***E***) Estimated maximum response strength as a function of age. ***F)*** Estimated decay time constant as a function of age.

### Qualitative features of LSO cells and their somatic inputs

An increase in apparent single fiber strength can have various presynaptic causes. One possible cause is the addition of new synaptic connections from each fiber. Alternatively, individual synapses could be strengthened through such morphological changes as increases in bouton or active zone size, or in number or distribution of synaptic vesicles. To distinguish among these possibilities, we analyzed the ultrastructure of synaptic inputs to the somata of LSO neurons at P1/2, P6/7, and P12/13 using electron microscopy (EM) (Fig. 2). Principal neurons of the medial and middle limbs of the LSO (n = 6 cells per age; N = 2 animals per age; Fig. 2A) were identified in the electron micrographs by their size, fusiform shape, and orientation with respect to the principal axis of the LSO (Helfert and Schwartz, 1987; Rietzel and Friauf, 1998). Synaptic boutons were identified by shape, proximity to soma, and presence of vesicles. As vesicles were sparse at the youngest ages, synapse-like structures that exhibited closely apposed membranes but that contained no vesicles in the plane of section were included in the analysis.

**Figure 2.**
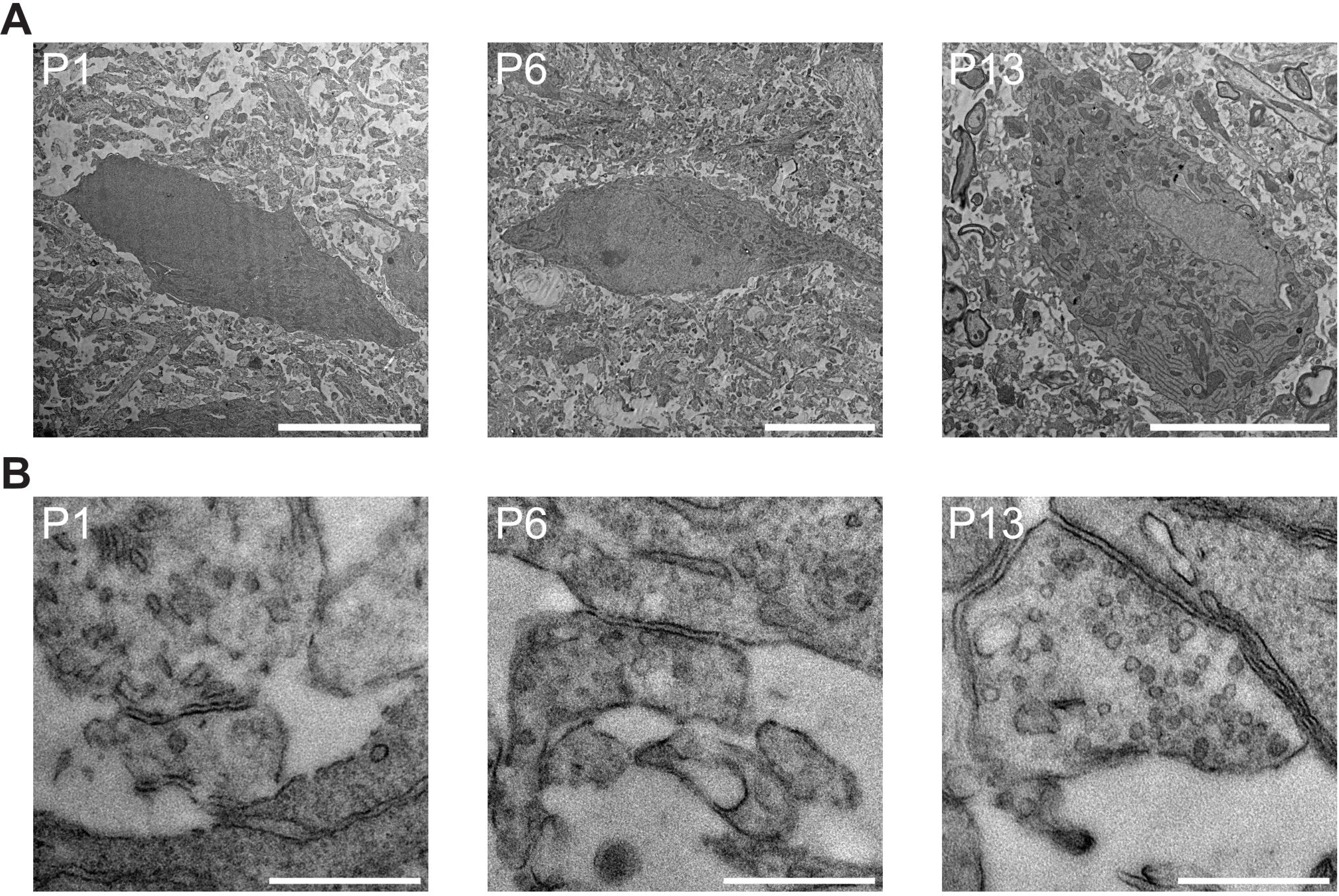
Representative TEM images of LSO neurons and their somatic synaptic inputs at P1, P6, and P13 ***A)*** Representative images of LSO neurons. Principal neurons were identified based on shape of cell body and orientation within the nucleus (n = 6 cells per each age group). Scale bars = 10 µm. ***B)*** Representative images of synaptic inputs to LSO soma. Images represent median vesicle count and median bouton size. Scale bars = 500 nm.

The representative images of Figure 2 illustrate some striking differences with respect to mature LSO tissue, as well as noticeable maturation during the first two postnatal weeks. Despite the demonstration of canonical excitatory inputs onto the somata of immature LSO principal cells at slightly older (P18) ages (Gjoni et al., 2018), we did not observe distinct postsynaptic densities (PSDs) at any of the ages examined here (Fig. 2B). In general, bouton size appeared to increase with age. Mitochondria were rare in the first week, appearing more reliably only in P12/13 boutons (Fig. 2B). No dense-core vesicles were present in any of the synaptic boutons we studied at any age. Relative to mature LSO tissue (Helfert et al., 1992), synaptic vesicles were sparse and loosely clustered at P12/13. Strikingly, at younger ages, synaptic vesicles were yet sparser or even absent at synaptic-like contacts (Extended Data Fig. 6-1A). Indeed, in tissue from the first week, one-third of synapse-like contacts onto the postsynaptic cell contained no apparent vesicles. For a closer analysis of each visually identified synapse, we used a modified ImageJ/FIJI macro (Extended Data Fig. 2-1) from SynapsEM (Watanabe et al., 2020) to manually annotate plasma membrane, mitochondria, putative active zones, and clear vesicles. As PSDs were not evident in any of these young synapses, putative active zones were defined as those areas where the membranes of the presynaptic and postsynaptic neurons were parallel and in close apposition (<20 nm separation).

### Bouton and mitochondrion size and number increase in second postnatal week

Given the importance of number and structure of boutons and mitochondria for synaptic transmission, we measured bouton area, active zone length, and mitochondrion area and number for our three age groups (Fig. 3). The number of putative boutons per soma in our sample did not change appreciably as a function of age (medians = 25, 22, 20.5 boutons; n = 6 cells each age; Fig. 3A). Bouton size increased little between P1/2 (median = 0.18 µm^2^; n = 154) and P6/7 (median = 0.23 µm^2^; n = 146) but significantly by P12/13 (median = 0.63 µm^2^, n = 123; Fig. 3B-C). Putative active zones became longer with age (medians = 0.20, 0.24, and 0.31 µm; Fig. 3D; Extended Data Figure 3-1). Like bouton size, mitochondrion size increased most appreciably in the second postnatal week (medians = 0.04, 0.04, and 0.10 μm^2^; n = 32, 29, 112; Fig. 4A-C). Mitochondrion number also increased with age: less than 20% of boutons in P1-7 tissue contained any mitochondria, whereas 50% of P12/13 boutons contained mitochondria (Fig. 4A,C). Moreover, over 20% of boutons at P12/13 had multiple mitochondria (Fig. 4A,C). These results suggest that bouton growth and recruitment or fission of mitochondria occur after the first postnatal week.

**Figure 3.**
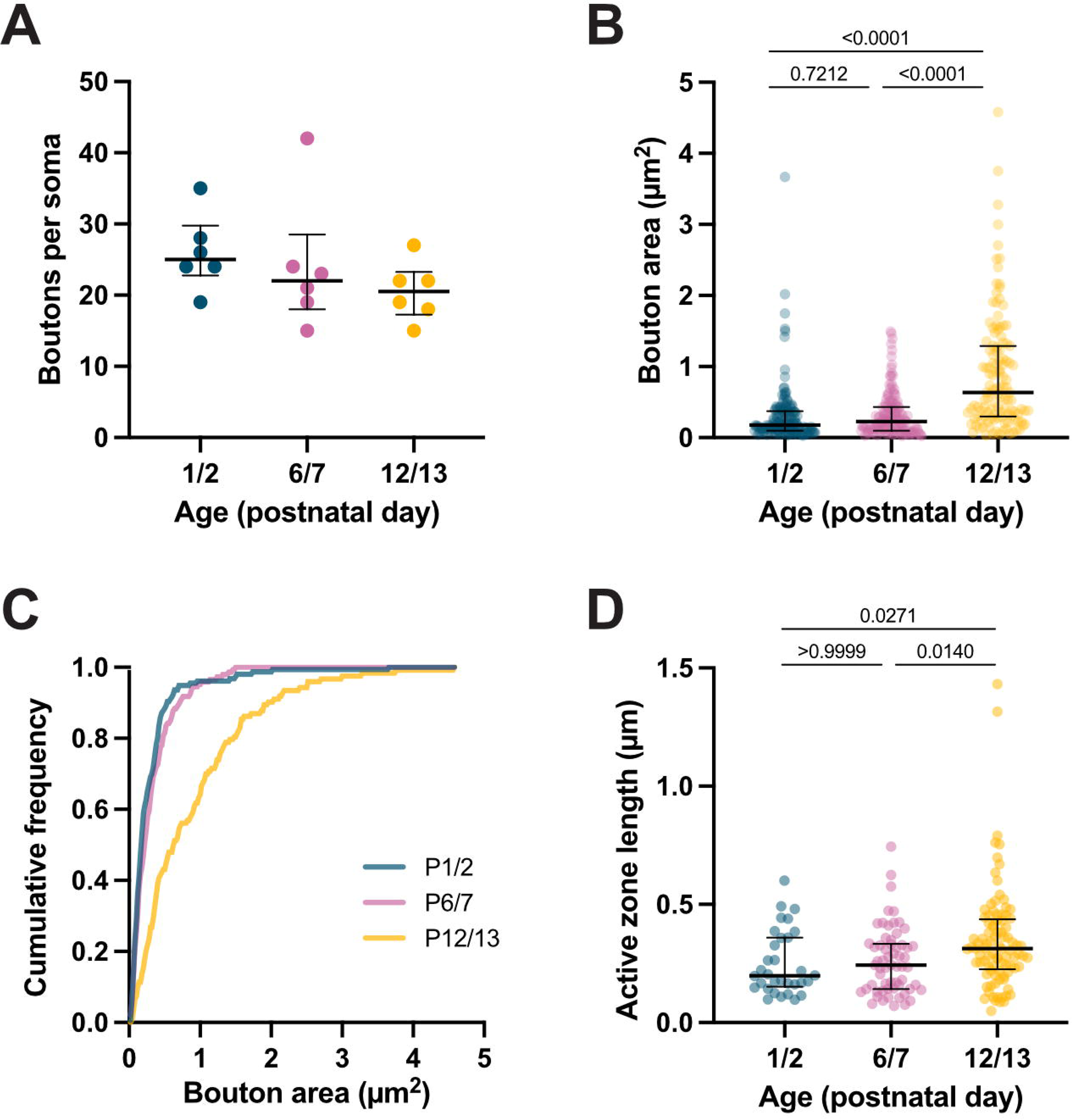
Bouton size and active zone length increased between P1 and P13. ***A)*** Number of boutons per soma between P1 and P13. Bouton counts (rounded to the nearest integer) for 25% quartile/***Median***/75% quartile were: 23/25/30 at P1/2 (blue); 18/22/29 at P6/7 (pink), and 17/21/23 at P12/13 (yellow) (n = 6 cells per age). ***B)*** Bouton area at P1/2, P6/7, and P12/13. Values (in µm^2^) for 25% quartile/***Median***/75% quartile were: 0.10/***0.18***/0.37 at P1/2 (n = 154); 0.09/***0.22***/0.43 at P6/7 (n = 146); 0.30/***0.63***/1.3 at P12/13 (n = 123); (p < 0.0001, Kruskal-Wallis Test; P1/2 vs. P6/7, p = 0.0.7212; P1/2 vs. P12/13, p < 0.0001; P6/7 vs P12/13, p < 0.0001, Dunn’s Test). ***C)*** Cumulative distribution of bouton area at P1/2, P6/7, and P12/13. ***D)*** Active zone lengths at P1/2, P6/7, and P12/13. Values (in µm) were 25% quartile/***Median***/75% quartile (µm): 0.15/***0.20***/0.36 at P1/2 (n = 32); 0.14/***0.24***/0.33 at P6/7 (n = 61); 0.23/***0.31***/0.43 at P12/13 (n = 93); (p < 0.0001, Kruskal-Wallis Test; P1/2 vs. P6/7, p > 0.9999; P1/2 vs. P12/13, p = 0.0271; P6/7 vs P12/13, p < 0.0140, Dunn’s Test).

**Figure 4.**
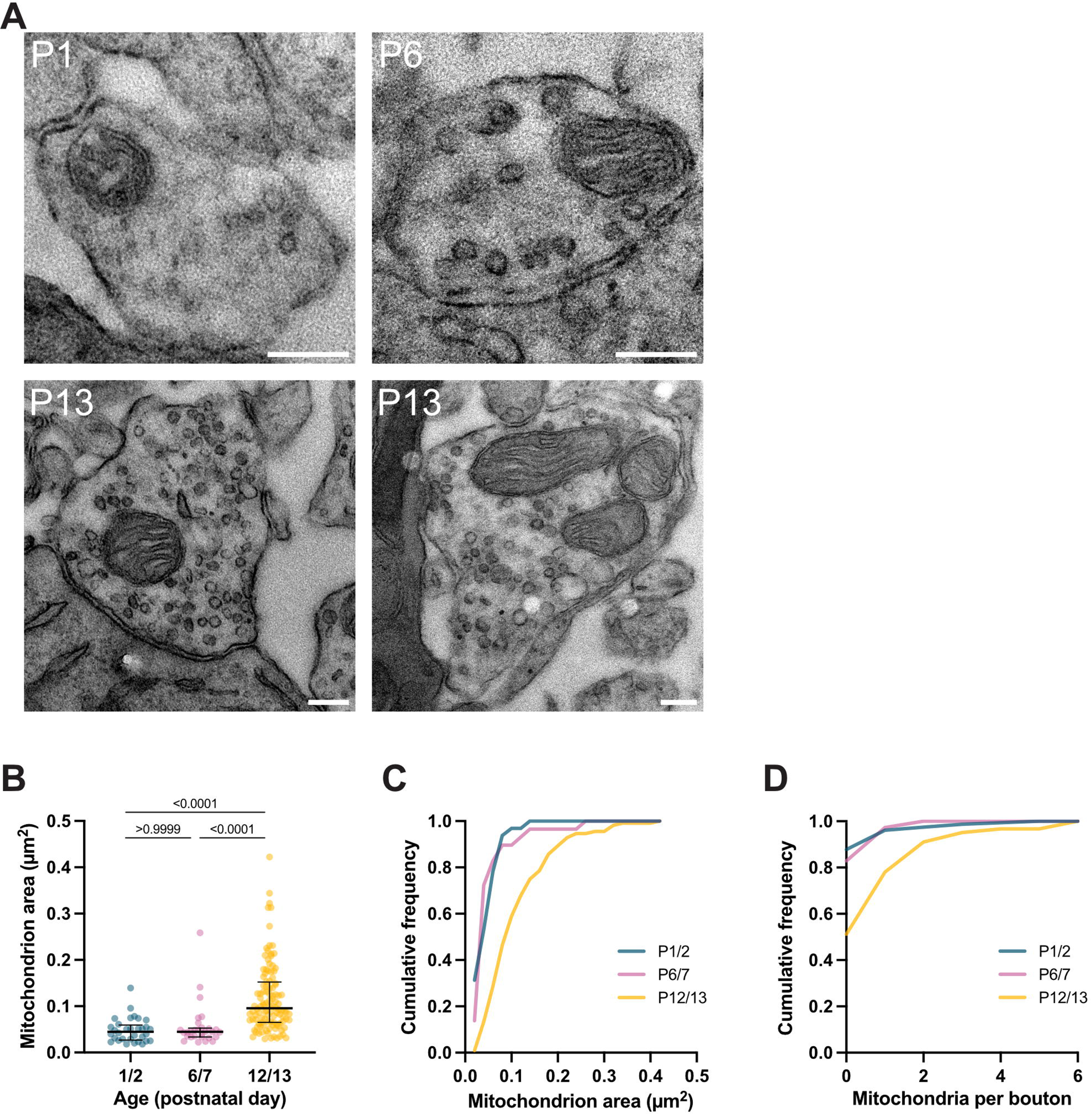
Mitochondrion size and number in boutons increased between P1 and P13. ***A)*** Examples of boutons with mitochondria between P1 and P13. ***B)*** Mitochondrion size at ages P1/2, P6/7, and P12/13. Values (in µm^2^) were 25% quartile/***Median***/75% quartile: 0.03/***0.04***/0.06 at P1/2 (n = 32); 0.03/***0.04***/0.05 at P6/7 (n = 29); 0.07/***0.1***/0.15 at P12/13 (n = 112); (p < 0.0001, Kruskal-Wallis Test; P1/2 vs. P6/7, p > 0.9999; P1/2 vs. P12/13, p < 0.0001; P6/7 vs P12/13, p < 0.0001, Dunn’s Test). ***C)*** Cumulative distribution of mitochondrion area at P1/2, P6/7, and P12/13. ***D)*** Cumulative mitochondrion frequency per bouton. Values for 25% quartile/***Median***/75% quartile were: 0/***0***/0 at P1/2 (n = 154); 0/***0***/0 at P6/7 (n = 146); 0/***0***/1 at P12/13 (n = 123); (p < 0.0001, Kruskal-Wallis Test; P1/2 vs. P6/7, p > 0.9999; P1/2 vs. P12/13, p < 0.0001; P6/7 vs P12/13, p < 0.0001, Dunn’s Test).

### Synaptic vesicle dimensions

To determine whether the period of major pre-hearing functional refinement in the LSO is accompanied by physical changes in vesicles, we measured the area and shape of all vesicles in the dataset (Fig. 5). At all ages, >70% of all vesicles had cross-sectional areas that fell in the range 1200-2400 nm^2^ (Fig. 5A), and no changes in vesicle size were observed with age. We also measured the major and minor axes to compute an aspect ratio (AR; major/minor axis) for each vesicle (Fig. 5B-D). At all ages examined, ARs for the entire vesicle population were similarly distributed: most vesicles were elliptical, and the distribution of AR was skewed rightward, with over half of all vesicles exhibiting an AR >1.2 (Fig. 5C-D). The large variability in vesicle shape across all synapses was consistent with an expected mix of excitatory and co-transmitting or inhibitory inputs at these ages (Gillespie et al., 2005; Gjoni et al., 2018).

**Figure 5.**
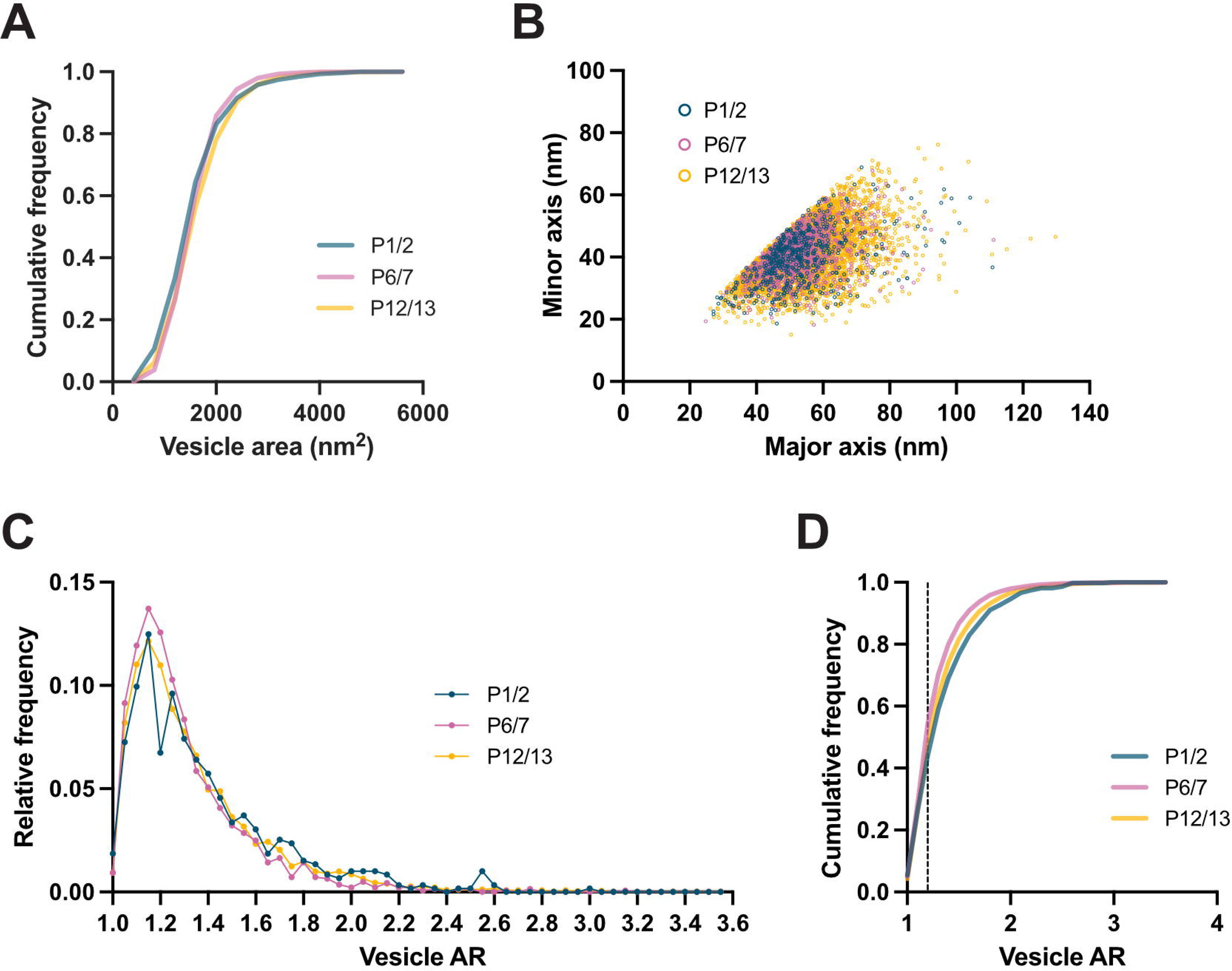
Synaptic vesicle dimensions were consistent between P1 and P13. ***A)*** Frequency distribution of vesicle size at P1/2, P6/7, and P12/13. Values for 25% quartile/***Median***/75% quartile (in nm^2^) were: 1283/***1617***/ 2013 at P1/2 (n = 572; blue); 1384/***1683***/1981 at P6/7 (n = 1401; pink); 1371/***1706***/2126 at P12/13 (n = 5148; yellow) (p < 0.0001, Kruskal-Wallis Test; P1/2 vs. P6/7, p = 0.0257, P1/2 vs. P1213, p < 0.0001, P6/7 vs. P12/13, p < 0.0184, Dunn’s Test). ***B)*** Scatter plot showing major and minor axes for all vesicles at P1/2, P6/7, P12/13. ***C)*** Frequency distribution of vesicle AR at P1/2, P6/7, P12/13. ***D)*** Cumulative distribution of vesicle AR at P1/2, P6/7, P12/13. Dashed line indicates AR = 1.2.

### Synaptic vesicle number and distribution change with age

Vesicle number and distribution varied with age over the first two postnatal weeks (Fig. 6-7). Median vesicle number within the bouton increased by a factor of 2 between P1/2 and P6/7, and more dramatically between P6/7 and P12/13, by a factor of nearly 8 (medians = 2, 4, and 30 vesicles; n = 154, 146, 123 synaptic profiles; Fig. 6A; Extended Data 6-1). At the two earliest ages, a large fraction of boutons (60% and 40%) had fewer than 5 vesicles, whereas over 90% of P12/13 boutons had at least 5 vesicles (Fig. 6B; Extended Data 6-1). Despite the growth in bouton size with age, vesicle density increased, roughly doubling between each age-group tested (medians = 9, 20, and 42 vesicles/µm^2^; Fig. 6C-D). To examine spatial distribution of vesicles, we calculated the distances (center-to-center) of each vesicle to its 3 nearest-neighbor vesicles (Fig. 7A). The distances between nearest-neighbor vesicles decreased with age and were smaller at all ages than the average nearest-neighbor distances predicted by vesicle density (Fig. 7A), suggesting that vesicles are already somewhat clustered within the bouton at early postnatal ages. The number of vesicles within 1 µm of the active zone increased with age with relatively similar distribution of vesicles throughout the bouton between age groups (Fig. 7B-C). Without imposing assumptions about expected distances from the active zone for readily releasable, replacement, or recycling pools, we asked how many vesicles were found within 50 nm (medians = 0, 1, and 1 vesicle; Fig. 7D) and 100 nm of the active zone (medians = 1, 2, and 4 vesicles; Fig. 7E). These increases in vesicle number near the active zone are consistent with a steady increase in the availability of vesicles in the recycling or readily-releasable pool (Schikorski and Stevens, 2001).

**Figure 6.**
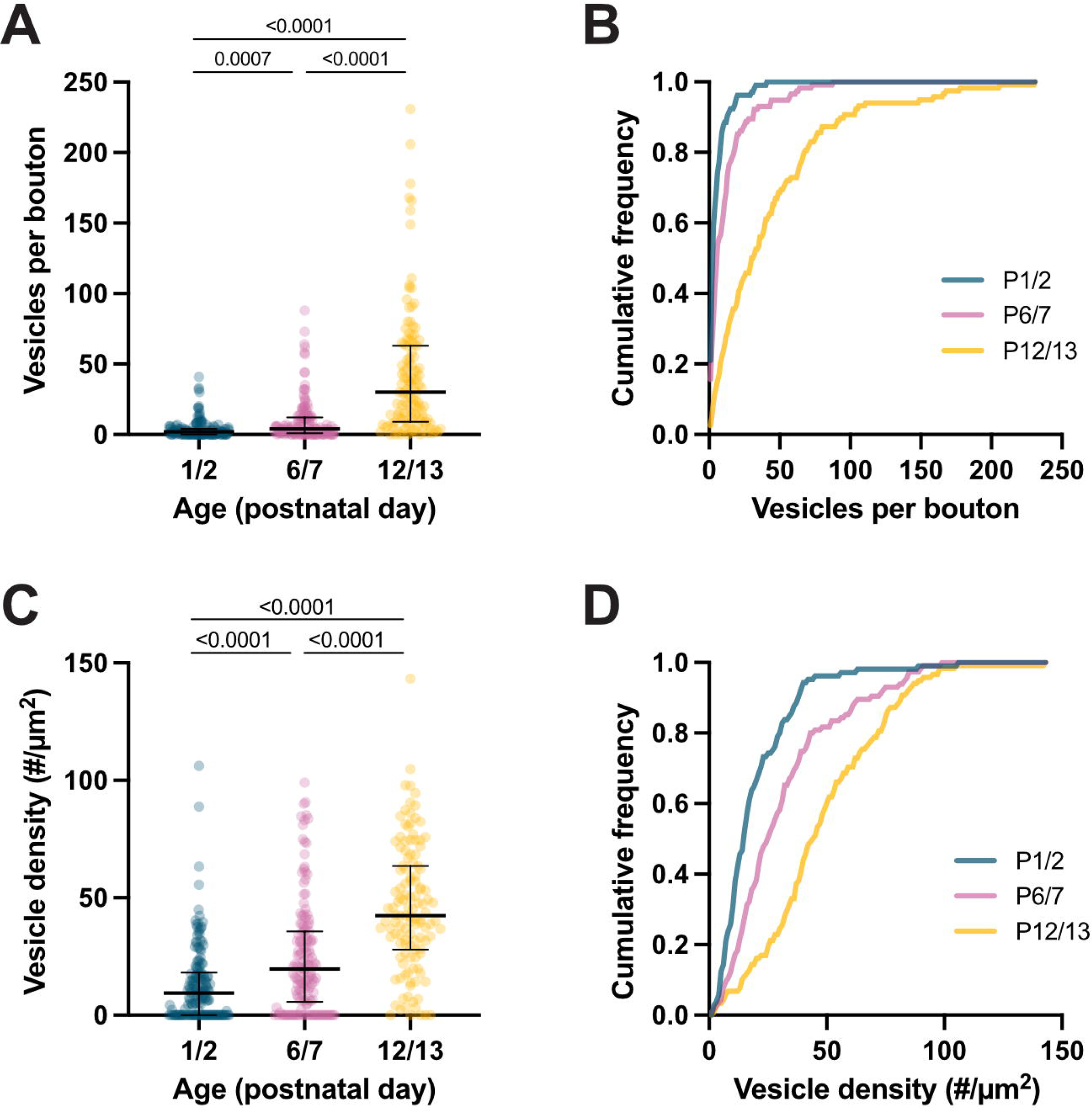
Number of vesicles per bouton increased between P1 and P13. ***A)*** Vesicle numbers per bouton at P1/2 (blue), P6/7 (pink), and P12/13 (yellow). Vesicle counts (rounded to nearest integer) for 25% quartile/***Median***/75% quartile were: 0/***2***/4 at P1/2 (n = 154); 1/***4***/12 at P6/7 (n = 146); 9/**30**/63 at P12/13 (n = 123); (p < 0.0001, Kruskal-Wallis Test; P1/2 vs. P6/7, p = 0.0007; P1/2 vs. P12/13, p < 0.0001; P6/7 vs P12/13, p < 0.0001, Dunn’s Test). ***B)*** Cumulative distribution of vesicle numbers per bouton at P1/2, P6/7, P12/13. ***C)*** Vesicle density within boutons at P1/2, P6/7, and P12/13. Values (count/µm^2^; rounded to nearest integer) for 25% quartile/***Median***/75% quartile were: 0/***9***/18 P1/2 (n = 154); 5.75/***19.72***/35.65 P6/7 (n = 146); 27.94/***42.45***/63.61 P12/13 (n = 123); (p < 0.0001, Kruskal-Wallis Test; P1/2 vs. P6/7, p < 0.0001; P1/2 vs. P12/13, p < 0.0001; P6/7 vs P12/13, p < 0.0001, Dunn’s Test). ***D)*** Cumulative distribution of vesicle density at P1/2, P6/7, P12/13.

**Figure 7.**
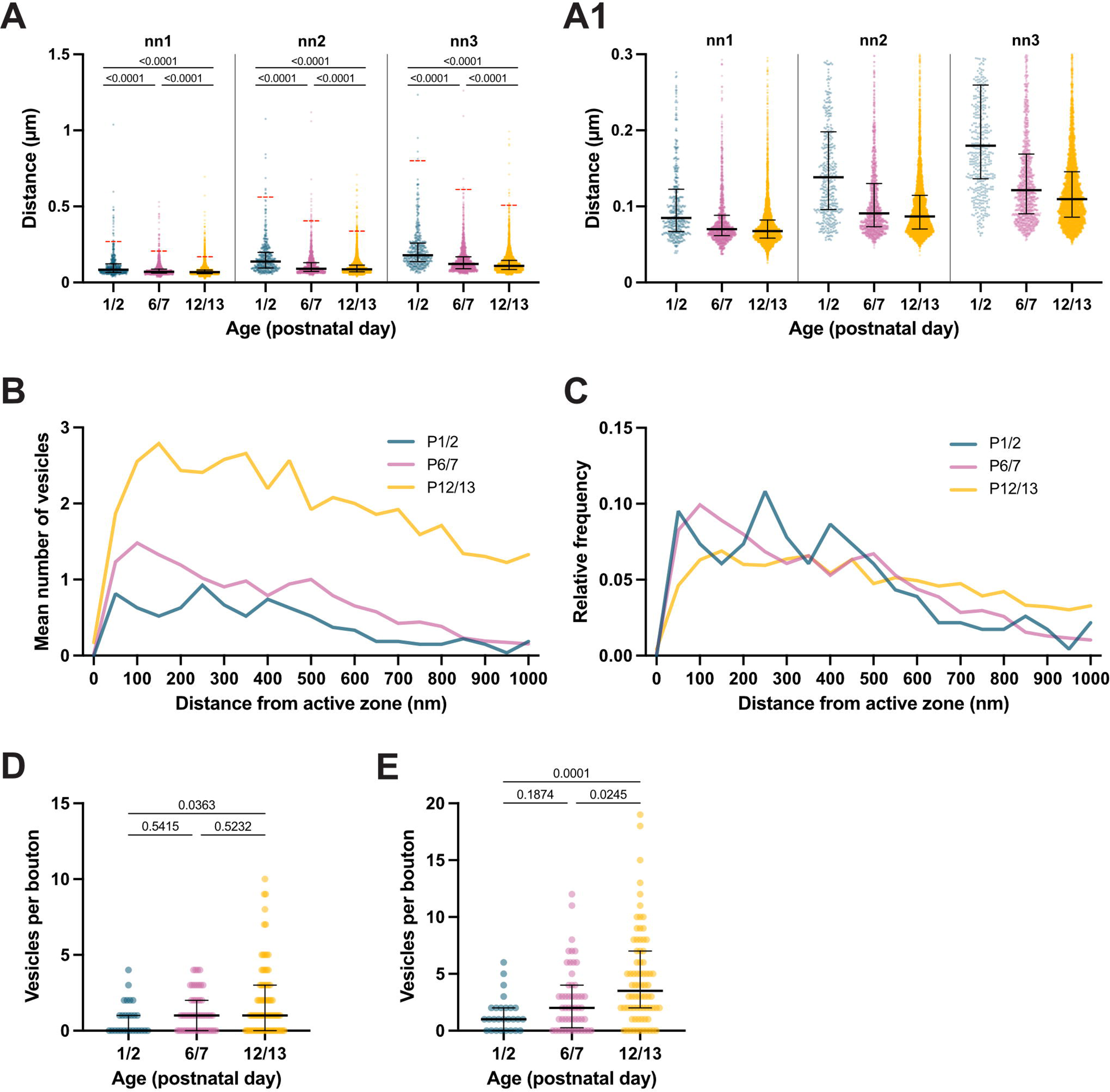
Spatial distribution of vesicles in presynaptic inputs to LSO neurons from P1 to P13 ***A*)** Distance between vesicles decreased between P1 and P13. Values (in nm) for 25% quartile/***Median***/75% quartile of first nearest neighbor (nn1) were: 67.13/***84.85***/122.50 at P1/2 (n = 450; blue); 61.55/***70.19***/88.29 at P6/7 (n = 1293; pink); 58.35/***67.56***/82.04 P12/13 (n = 5112; yellow); (p < 0.0001, Kruskal-Wallis Test; P1/2 vs. P6/7, p < 0.0001; P1/2 vs. P12/13, p < 0.0001; P6/7 vs P12/13, p < 0.0001, Dunn’s Test). Values (in nm) for 25% quartile/***Median***/75% quartile of second nearest neighbor (nn2) were: 95.81/***138.50***/198.10 at P1/2; 73.42/***90.87***/130.20 at P6/7; 70.38/***86.96***/114.5 at P12/13; (p < 0.0001, Kruskal-Wallis Test; P1/2 vs. P6/7, p < 0.0001; P1/2 vs. P12/13, p < 0.0001; P6/7 vs P12/13, p < 0.0001, Dunn’s Test). Values (in nm) for 25% quartile/***Median***/75% quartile of third nearest neighbor (nn3) were: 136.50/***179.90***/259.60 at P1/2; 90.36/***125.50***/168.90 at P6/7; 85.86/***109.70***/145.6 P12/13; (p < 0.0001, Kruskal-Wallis Test; P1/2 vs. P6/7, p < 0.0001; P1/2 vs. P12/13, p < 0.0001; P6/7 vs P12/13, p < 0.0001, Dunn’s Test). Dotted red line represents nearest neighbor vesicle distance given vesicle density with uniform vesicle distribution. ***A1*)** Same data as in ***A***, showing nearest neighbor distances within 300 nm. ***B)*** Mean vesicle number as function of distance from active zone. ***C)*** Normalized distribution of vesicle distance from active zone. ***D)*** Number of vesicles within 50 nm of active zone increases with age. Values (count) for 25% quartile/***Median***/75% quartile were: 0/***0***/1 at P1/2 (n = 27); 0/***1***/2 at P6/7 (n = 52); 0/***1***/3 at P12/13 (n = 76); (p < 0.0001, Kruskal-Wallis Test; P1/2 vs. P6/7, p < 0.5415; P1/2 vs. P12/13, p < 0.0363; P6/7 vs P12/13, p < 0.5232, Dunn’s Test). ***E)*** Number of vesicles within 100 nm of active zone increases with age Values (count; rounded to nearest integer) for 25% quartile/***Median***/75% quartile were: 0/***1***/2 at P1/2 (n = 27); 0/***2***/4 at P6/7 (n = 52); 2/***4***/7 at P12/13 (n = 76); (p < 0.0001, Kruskal-Wallis Test; P1/2 vs. P6/7, p < 0.1874; P1/2 vs. P12/13, p < 0.0001; P6/7 vs P12/13, p < 0.0245, Dunn’s Test).

### Relative proportion of excitatory and inhibitory inputs changes with age

The aspiny nature of LSO neurons, together with the immature synapse features observed here (e.g., lack of PSDs, sparsity of vesicles), introduced additional uncertainty to using solely ultrastructural features to categorize immature synapses as excitatory or inhibitory. Therefore, we first estimated the relative proportions of excitatory and inhibitory somatic synapses by labeling synaptic proteins. In a separate set of tissue (2 animals per age), we co-immunostained for synapsin, vesicular glutamate transporter 1 (VGLUT1), and vesicular inhibitory amino acid transporter (VIAAT), and then imaged with a confocal microscope (Fig. 8). We used synapsin immunoreactivity to select synapses around LSO somata (n = 19, 21, 23 cells), and categorized the selected synapses as excitatory if they were VGLUT1^+^ or inhibitory if they were VIAAT^+^ within a 200 nm search radius (Fig. 8A-B). This searching method allows for small, nearby synapses to register as overlapping, and thus, to estimate uncertainty, we tested what fraction of VGLUT1+ terminals overlapped with VIAAT, and vice versa (VGLUT1 fraction VIAAT^+^: 8%, 13%, 8%; VIAAT fraction VGLUT1^+^: 8%, 8%, 4%). At each age, a significant proportion of synapses could not be categorized, likely because of our stringent (≥4 S.D.) criterion for synapse assignment (Fig. 8C). At P1, most somatic synapses were excitatory (52% VGLUT1-IR vs. 21% VIAAT-IR). By P6, excitatory synapses were less common around the soma (23% VGLUT1-IR vs. 47% VIAAT-IR), and by P12, excitatory synapses were even less common at the soma (13% VGLUT1-IR vs. 46% VIAAT-IR) (Fig. 8C). Thus, the general trend at LSO somata during the pre-hearing postnatal period was for the fraction of excitatory boutons to decrease and the fraction of inhibitory boutons to increase.

**Figure 8.**
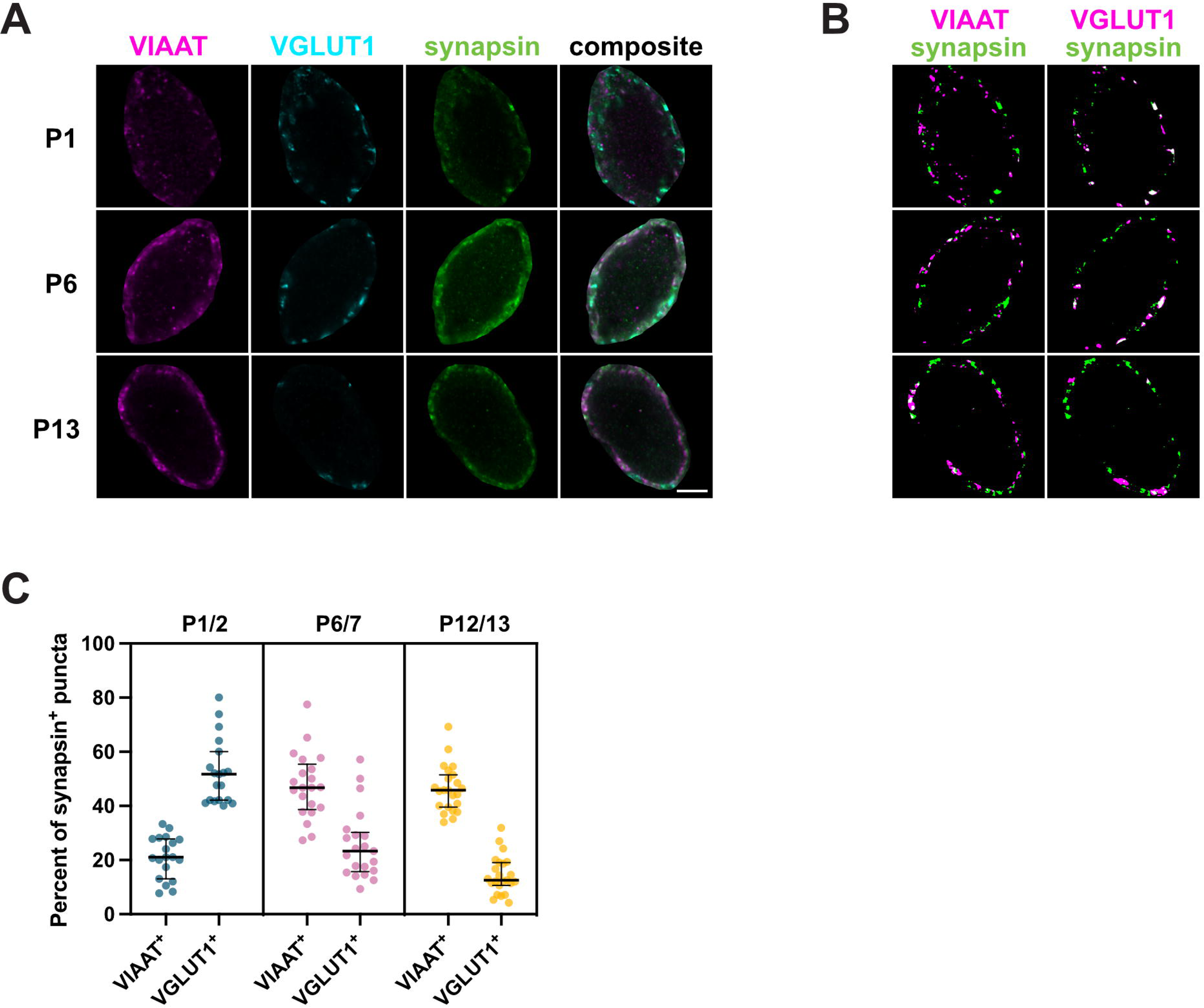
Relative fraction of excitatory synapses declines, and fraction of inhibitory synapses increases, for inputs to LSO somata, between P1 and P13. ***A)*** Representative LSO soma showing VIAAT-(magenta), VGLUT1-(cyan), and synapsin-(green) immunoreactivity (IR) at P1, P6 and P13. Scale bar = 2 µm. ***B)*** Thresholded images of representative LSO soma showing overlap of synapsin (green) and VIAAT (magenta) on ***left*** and overlap of synapsin (green) and VGLUT1 (magenta) on ***right***. ***C)*** Fraction of total synapsin^+^ puncta that overlap with VIAAT-IR and VGLUT1-IR at P1/2, P6/7, and P12/13. Values (in %) for 25% quartile/***Median***/75% quartile for synapsin-IR and VIAAT-IR overlap were: 13.04/***21.05***/27.78 at P1/2 (n = 19); 34.59/***46.67***/55.36 at P6/7 (n = 21); 39.58/***45.83***/51.43 at P12/13 (n = 23). Values (in %) for 25% quartile/***Median***/75% quartile for synapsin-IR and VGLUT1-IR overlap were: 42.11/***51.72***/60.00 at P1/2 (n = 19); 15.69/***23.33***/30.21 at P6/7 (n = 21); 10.61/***12.50***/19.05% at P12/13 (n = 23).

We then turned to the ultrastructural data set to attempt to categorize synapses according to vesicle shape. Based on well-established characteristics of mature excitatory and inhibitory synapses, we expected an excitatory synapse to contain mostly spherical vesicles and an inhibitory synapse to contain mostly ellipsoidal vesicles (Gray, 1969; Tao et al., 2018). Simple co-transmitting synapses, in which GABA/glycine and glutamate are released from separate vesicles, might then contain a mix of spherical and ellipsoidal vesicles. Under this model, excitatory synapses should contain spherical vesicles, with mean vesicle AR near 1. Inhibitory synapses should have a larger mean vesicle AR as well as an AR variance made larger by the process of randomly sectioning ellipsoids. Ignoring P1/2 boutons, which had too few vesicles for analysis, we calculated the mean AR for every bouton containing 5 or more vesicles at P6/7 (n = 67) and at P12/13 (n = 104).

To assess bouton phenotype using average AR, we examined P12/13 boutons, given that most synapses at this age are expected to be either excitatory or inhibitory, whereas many synapses at P6/7 may be co-transmitting. The distribution of mean ARs for P12/13 boutons revealed two local maxima that could reflect mean values for excitatory (1.25) and inhibitory (1.44) populations. Intriguingly, the distribution for P6/7 was also bimodal, with a first peak (1.42) close to the value of the larger P12/13 peak, and a second peak whose value (1.30) lay between the two P12/13 maxima. We used the value of the trough between the two P12/13 peaks (1.33) to sort synapses into putative excitatory (mean AR < 1.33) and putative inhibitory (mean AR > 1.33) synapses (Fig. 9A). Putative excitatory synapses appeared to contain mostly round vesicles whereas putative inhibitory synapses appeared to contain a mixture of round and elliptical vesicles (Fig. 9B). We did not observe any differences in PSD thickness between putative excitatory and inhibitory synapses. We then plotted the ARs for all vesicles within these categories at P6/7 (n = 987 vesicles from putative excitatory synapses, 306 vesicles from putative inhibitory synapses; Fig. 8C) and P12/13 (n = 2391 vesicles from putative excitatory synapses, 2721 vesicles from putative inhibitory synapses; Fig. 9D). As expected, ARs were more variable for the putative inhibitory vesicle populations (Full width half maximum = 0.28 excitatory vs 0.38 inhibitory for P6/7; 0.29 excitatory vs 0.50 inhibitory for P12/13). Using this average AR criterion to categorize terminal type, we asked what proportions of identified synapses onto each cell were putative excitatory and putative inhibitory at P6/7 and at 12/13 (Fig. 9E). These fractions differ from those found by immunolabeling for synaptic proteins. Nevertheless, regardless of analysis, the general trend held for somatic synapses: at early ages excitatory synapses were most abundant, and by hearing onset inhibitory synapses were most abundant. Thus, even at immature synapses that lack classical features of mature synapses, vesicle shape may be useful in identifying synapse type. Visual inspection of P6/7 boutons yielded no indications of separate vesicle pools in presumed co-transmitting synapses.

**Figure 9.**
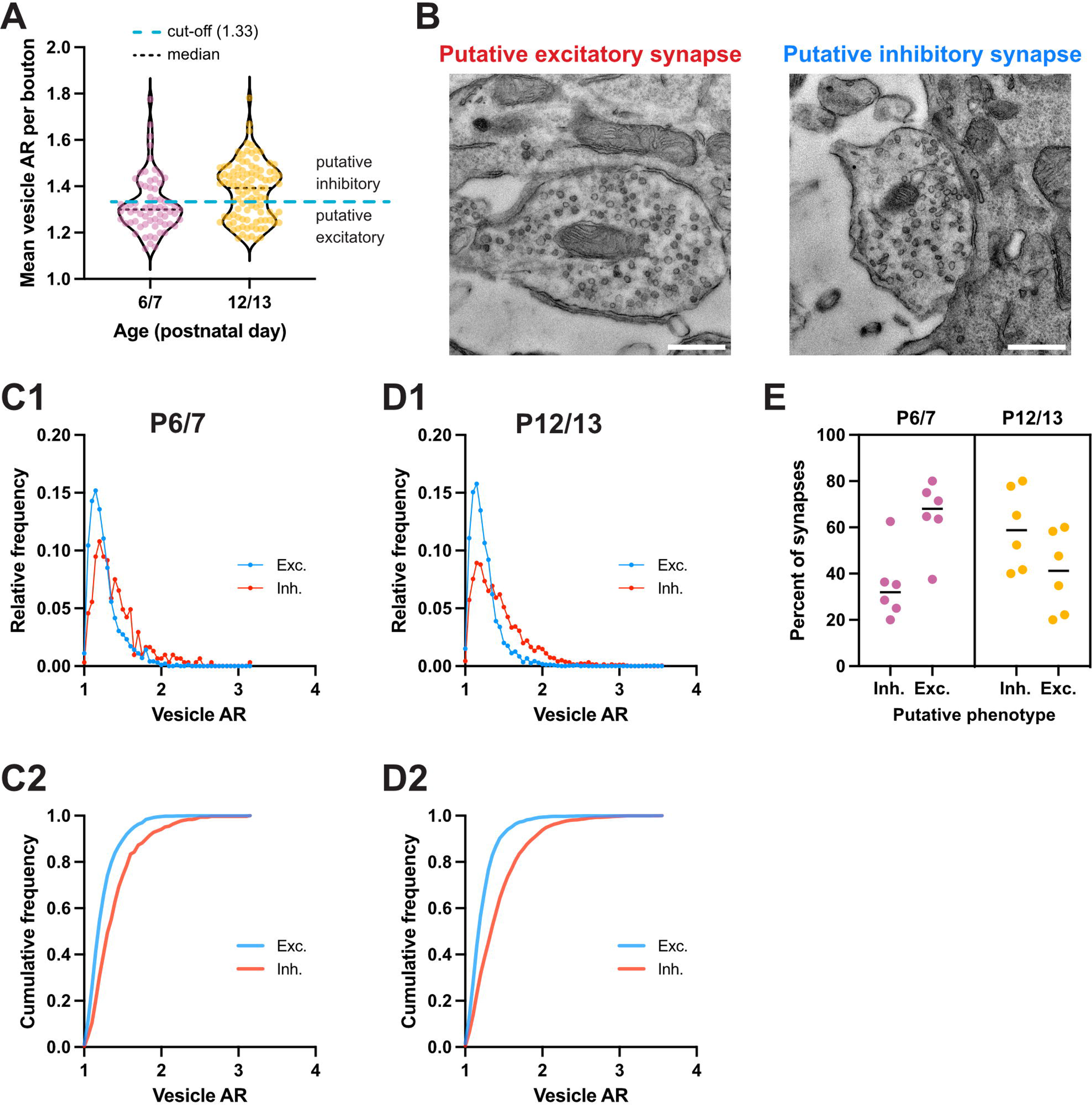
Identification of synapse type based on vesicle shape ***A)*** Mean vesicle AR per bouton for all P6/7 (n = 67; pink) and P12/13 (n = 104; yellow) synapses with ≥ 5 vesicles. Putative inhibitory and putative excitatory synapses were partitioned based on the bouton’s mean AR, using a partition value (light blue) of 1.33. ***B)*** Example of a putative excitatory and putative inhibitory synapse. Scale bar = 500 nm. ***C1-2)*** Relative and cumulative frequency distribution of vesicle ARs in putative excitatory and inhibitory synapses for P6/7 (n = 987 vesicles for excitatory synapses, 306 vesicles for inhibitory synapses). Full width at half maximum (FWHM) = 0.28 for excitatory synapses, 0.38 for inhibitory synapses. ***D1-2)*** Relative and cumulative frequency distribution of vesicle ARs in putative excitatory and inhibitory synapses for P12/13 (n = 2391 vesicles for excitatory synapses, 2721 vesicles for inhibitory synapses). FWHM = 0.29 for excitatory synapses, 0.50 for inhibitory synapses. ***E)*** Relative percentage of putative inhibitory and excitatory synapses for each cell at P6/7 and P12/13 (n = 6 cells per age).

Having identified putative excitatory and inhibitory synapses using the average vesicle AR, we re-examined the dataset to look for any surface-level features that might allow us to distinguish synaptic phenotypes. After assigning the synapses to an excitatory or inhibitory phenotype based on vesicle shape, we were unable to detect any differences in vesicle, bouton, or mitochondrion parameters between excitatory and inhibitory synapses (data not shown). These observations suggest that these excitatory and inhibitory synapses mature similarly and according to similar time courses.

## Discussion

Our goal was to characterize the ultrastructure of immature synaptic inputs to LSO cell somata and to discover whether or what structural changes at individual synapses accompany pre-hearing functional refinement of these inputs. Although the AVCN and MNTB make functional connections onto LSO principal cells before birth (Kandler and Friauf, 1995), even at birth these synapses had a strikingly immature appearance. Subsequently, somatic LSO synapses exhibited increases in bouton size, vesicle number, and in the number and size of mitochondrion, all during a pre-hearing period that correlates with the period of major functional refinement. Our results support a model in which, in addition to an increase in synapse number, physical changes at presynaptic terminals contribute to functional strengthening.

### Maturation of synaptic inputs to the LSO before hearing onset

The primary excitatory and inhibitory inputs to principal cells of the LSO originate in the ipsilateral VCN and MNTB (Glendenning et al., 1985), though LSO cells can also receive inhibitory inputs from other sources (Gomez-Alvarez and Saldana, 2016; Jalabi et al., 2013, Kuwabara and Zook, 1992; Warr and Beck, 1996; Weingarten et al., 2023); hence, we make the simplifying assumption that most of the synapses studied here represent excitatory terminals from the AVCN and inhibitory terminals from the MNTB. As a population, synaptic terminals in the LSO exhibited striking changes between birth and hearing onset, with vesicle number increasing by a factor of 15 and vesicle density by a factor of 5. Interestingly, despite a significant increase in functional input size in the first week (Kim and Kandler, 2003; Kim and Kandler, 2010), bouton and mitochondrion size remained largely unchanged until the following week, when they increased by factors of 3 and 2, suggesting that these features are not primarily responsible for single-fiber strengthening in the first week. By P13, roughly half of all boutons sampled contained at least one mitochondrion. The increases in vesicle number in the first postnatal week may have supported, in part, the increases in input strength seen between P1 and P6, whereas later changes in bouton size and mitochondrion parameters could underlie other physiological maturation. Although refinement measured in the living slice appears to be largely complete by hearing onset (Walcher et al., 2011), synapses onto LSO somata at hearing onset had a much less mature appearance than those from juvenile and adult animals (Cant, 1984; Gjoni et al., 2018; Helfert et al., 1992). Additional changes after hearing onset, including vesicle number and bouton size, may further strengthen individual synapses, or perhaps help to sustain rapid synaptic transmission characteristic of auditory synapses (Brill et al., 2019).

In addition to changes in synapse structure, inputs to the LSO soma also undergo redistribution before hearing onset. Whereas somatic excitatory synapses outnumber inhibitory synapses at birth, somatic excitatory synapses have been lost and inhibitory synapses outnumber excitatory synapses by hearing onset. In the neighboring medial superior olive (MSO) nucleus of gerbil, inhibitory synapses are redistributed toward the soma after hearing onset, under the direction of acoustic activity (Kapfer et al., 2002; Magnusson et al., 2005). The two nuclei are not directly comparable, due to species differences as well as circuit organization (Grothe et al., 2010).

Nevertheless, MNTB axons undergo further rearrangement in the LSO in the 1-2 weeks after hearing onset (Sanes and Siverls, 1991), and synapses in the rat LSO experience further plasticity that may, as in the MSO, be influenced by acoustic activity.

Classically, excitatory synapses are defined in TEM by the presence of spherical vesicles, apposition to a dendritic spine, and a thick PSD, whereas inhibitory synapses are defined by the presence of ellipsoid vesicles and a thin PSD (Gray, 1969). These features, gleaned from early studies of mature synapses, apply less well to immature synapses in the developing nervous system (Ahmari and Smith, 2002). Although LSO principal cells generally lack spines (Cant, 1984; Helfert and Schwartz, 1986), glutamatergic transmission from the VCN is detectable before birth (Kandler and Friauf, 1995), and so the presence of synapses without identifiable PSDs was surprising. Notably, the classification – based on vesicle shape – that is presented here is consistent with the trends of synapse redistribution seen with immunohistochemistry.

Before hearing onset, MNTB terminals in the LSO release glutamate and glycine/GABA. During this period of prominent glutamatergic transmission (Gillespie et al., 2005), perisomatic VGLUT3 expression is common (Blaesse et al., 2005; Cooper and Gillespie, 2011). As most somatic synapses in the adult are inhibitory (Cant, 1984; Friauf et al., 1997; Helfert et al., 1992), we propose that at P6/7, the population of somatic synapses includes a large fraction of co-transmitting synapses from the MNTB (VGLUT3^+^/VIAAT^+^), and smaller proportions of other inhibitory (VIAAT^+^) inputs and of excitatory (VGLUT1^+^) inputs. These co-transmitting synapses might contain vesicles with distinct vesicular transporters (simple co-transmission), or they might contain vesicles all of which express multiple vesicular transporters, resulting in co-release for each vesicle fusion event. Although this question has not been examined directly, differences in short-term synaptic plasticity for glutamate and GABA/glycine transmission at the same MNTB-LSO synapses are consistent with a model in which the neurotransmitters occupy different sets of vesicles (Case and Gillespie, 2011), as reported at other synapses (Boulland et al., 2009; Root et al., 2018). Under the classical assumption that vesicle shape indicates the neurotransmitter therein, we expected the mean AR of a co-transmitting synapse to fall between those of excitatory and inhibitory synapses regardless of whether the synapse uses simple co-transmission or co-release. Although the distribution of ARs for P12/13 exhibited two distinct maxima, the distribution for P6/7 terminals displayed a single peak between the two P12/13 maxima, which we speculate could reflect a population of simple co-transmitting synapses that contain two types of vesicles with different average ARs. Lacking any way to test this speculation directly, we hesitate to use vesicle shape alone as a metric for distinguishing co-transmitting synapses from excitatory and inhibitory synapses; indeed, generalizing this method to distinguish excitatory from inhibitory synapses for different areas and synapses may require empirical estimation of AR.

In this first look at immature LSO synapses, we used 2D TEM images, rather than reconstructing a smaller number of synapses in 3D, to sample a larger number of synapses. As this choice carries some risk of sampling bias, and consequent underestimation of synaptic components or overestimation of numbers of putative boutons at young ages, the numbers provided here can be estimates at best. The relative numbers, however, are striking: the very youngest synapses, though functional, appear to have little in common with mature synapses, and over the next two weeks, these synapses undergo significant maturation. In addition to the striking structural changes within individual boutons, the distribution of synapses at LSO somata shifts from predominantly excitatory to predominantly inhibitory inputs.

### Structural basis for functional strengthening of inputs to the LSO

Functional refinement in the pre-hearing rodent MNTB-LSO pathway has been well characterized: over half of all single-fiber inputs are lost, the remaining inputs are strengthened, evoked miniature responses become larger, and release probability decreases (Kim and Kandler, 2003; Kim and Kandler, 2010; Nabekura et al., 2004). Structural studies have focused on morphology of MNTB inputs after hearing onset (Gjoni et al., 2018; Sanes and Siverls, 1991), and the speculation that addition of synapses from a single MNTB fiber supports functional strengthening of the MNTB-LSO pathway had not been previously examined at the EM level. The increase in inhibitory synapse density at the soma could result from addition of release sites at the soma, or from redistribution of existing synapses toward the soma. We find that the density of inhibitory synapses at the soma increases during the period of major refinement, consistent with a model in which single-fiber strengthening results from the addition of proximal release sites. Major changes in the physical structure of the synapse may underlie changes in quantal content (Kim and Kandler, 2010). Similar changes at the level of individual boutons almost certainly apply to the VCN-LSO pathway, which exhibits early functional refinement to a lesser extent (Case et al., 2011; Felix and Magnusson, 2016; Gjoni et al., 2018).

Throughout the nervous system, the physical size of the synapse and its various components correlates with functional synapse strength (Cheetham et al., 2014; Cserep et al., 2018; Harris and Stevens, 1988; Holler et al., 2021; Holtmaat and Svoboda, 2009; Murthy et al., 2001). At the presynaptic site, synapse strength is influenced by multiple factors, including vesicle availability, neurotransmitter concentration, release probability, number and location of release zone(s), and mitochondrial function (Edwards, 2007; Kaeser and Regehr, 2017; Vos et al., 2010). The growth of boutons and mitochondria, together with increases in vesicle number and vesicle density, seen here between P1 and P13, correlate with increased single fiber strength and may also support functionally stronger synapses. At hippocampal synapses, release probability scales linearly with the size of the active zone (Holderith et al., 2012), and although addition of vesicles near the active zone generally correlates with increased release probability, mere proximity to the active zone does not make vesicles fusion-ready. Furthermore, synapse maturation at excitatory and inhibitory synapses in the early auditory system involves a decrease in release probability (Brenowitz and Trussell, 2001; Kim and Kandler, 2010), consistent with maturation of timing. The increase in vesicle numbers within 100 nm of the active zone may reflect an increase in recycling or readily releasable pools and may contribute to an increase in quantal content throughout development. The largescale changes we see – including growth of bouton size, increased numbers of vesicles and mitochondria, and longer active zones – could make synapses more resilient in the face of the sustained, rapid synaptic transmission characteristic of auditory circuits (Brill et al., 2019; Krächan et al, 2017).

### General characteristics of immature synapses

In this first report on synaptic ultrastructure in the immature LSO, we found significant changes in the two weeks between birth and hearing onset. Hearing onset clearly does not mark the end of the developmental period, as our P13 synapses differed noticeably from older synapses in the LSO (P18 and adult) and elsewhere (Gjoni et al., 2018; Harris and Weinberg, 2012; Helfert et al., 1992). The developmental trends we saw were consistent with those seen at other immature synapses, including increases in vesicle number and in mitochondrion size and number (Adinolfi, 1972; Armstrong-James and Johnson, 1970; Dufour et al., 2016; Jones and Cullen, 1979; Markus et al., 1987). Although vesicles shrink across development in some areas (Dufour et al., 2016; Markus et al., 1987), we saw no changes in vesicle size in the pre-hearing LSO. The time course of maturation at LSO somatic synapses noticeably lagged that in the upstream MNTB. For example, at the Calyx of Held in the MNTB, vesicles and PSDs are reliably present as early as P2, and the terminal grows to roughly its mature volume by the end of the first postnatal week (Hoffpauir et al., 2006; Holcomb et al., 2013), whereas in the LSO vesicles were still sparse, and PSDs still absent, at P13. In light of the differential expression of PSD components with varying developmental expression patterns (Petralia et al., 2005) it is perhaps unsurprising that PSDs appear at different times across multiple preparations, or that, synapse maturation varies across brain regions and may encompass several weeks (Adinolfi, 1972; Armstrong-James and Johnson, 1970; Dufour et al., 2016; Jones and Cullen, 1979; Markus et al., 1987).

Identifying immature synapses and their phenotypes can be challenging, as we lack a universally-defined set of elements required for a functional immature synapse (Ahmari and Smith, 2002). These challenges only increase for immature synapses that release multiple neurotransmitters. Although synaptic phenotype (excitatory/inhibitory) has long been assigned on the basis of synapse location, vesicle shape, and postsynaptic density, variability in these parameters with age and area reduces the universal validity of this approach (Klemann and Roubos, 2011). Even at synapses that conform to the classical distinctions between symmetric and asymmetric synapses, differences in vesicle shape may reveal less about native synapses and more about how different synapses interact with fixation (Tao et al., 2018). The most promising current methods for establishing ground truth carry costs: immunogold labelling and correlative light and electron microscopy (CLEM) can help with synapse identification, usually at the cost of precise ultrastructure (Root et al., 2018), whereas genetically encoded probes for specific proteins identify vesicles, but at the risk of altering native expression (Shu et al., 2011; Steinkellner et al., 2021). The correlation shown here between aspect ratio- and synaptic label-based assignation of synapse phenotype offers one way to categorize immature synapses, with the caveat that an average AR cut-off for the binary categorization may need to be determined empirically for different preparations.

The significant changes in immature synapses of the LSO between birth (this study) and maturity (Helfert et al., 1992) highlight key general questions concerning the minimum common synaptic elements required for functional transmission, features that distinguish mature from immature synapses, and above all: the mechanistic processes that support synapse development and plasticity. Answers to these questions will require concerted, multipronged approaches marrying protein expression, ultrastructure, and physiology.

## Supporting information

Extended Data for Figure 2

Extended Data for Figure 3

Extended Data for Figure 6

List of antibodies and stains used for immunohistochemistry

---

**Extended Data Figure 2-1.** Example of modified SynapsEM annotation at a P13 synapse. ***A)*** Unannotated synapse. ***B)*** Synaptic vesicles (orange), mitochondria (blue), plasma membranes (green), and active zones (pink) were annotated using freehand tools in ImageJ/FIJI. Scale bar = 500 nm.

**Extended Data Figure 3-1.** Examples of additional boutons with observable active zones at P1 ***(A)***, P6 ***(B)***, and P13 ***(C)***. Scale bars = 200 nm.

**Extended Data Figure 6-1.** Variability of vesicle numbers in presynaptic inputs to LSO somata between P1 and P13. ***A)*** Examples of boutons with no observable vesicles present at P1, P6, and P13. ***B)*** Examples of boutons with vesicle numbers around 50% percentile at P1 (1 vesicle), P6 (3 vesicles), and P13 (33 vesicles). ***C)*** Examples of boutons with vesicle numbers above 90% percentile at P1 (33 vesicles), P6 (88 vesicles), and P13 (206 vesicles). Scale bars = 200 nm.

**Extended Data Table 7-1.** List of antibodies and stains used for immunohistochemistry.

## Notes

### Competing Interest Statement

The authors have declared no competing interest.

