## Supplementary figures and images for "Ultrastructure of immature synaptic inputs in the lateral superior olive of rodent brainstem"

### Extended Data for Figure 2

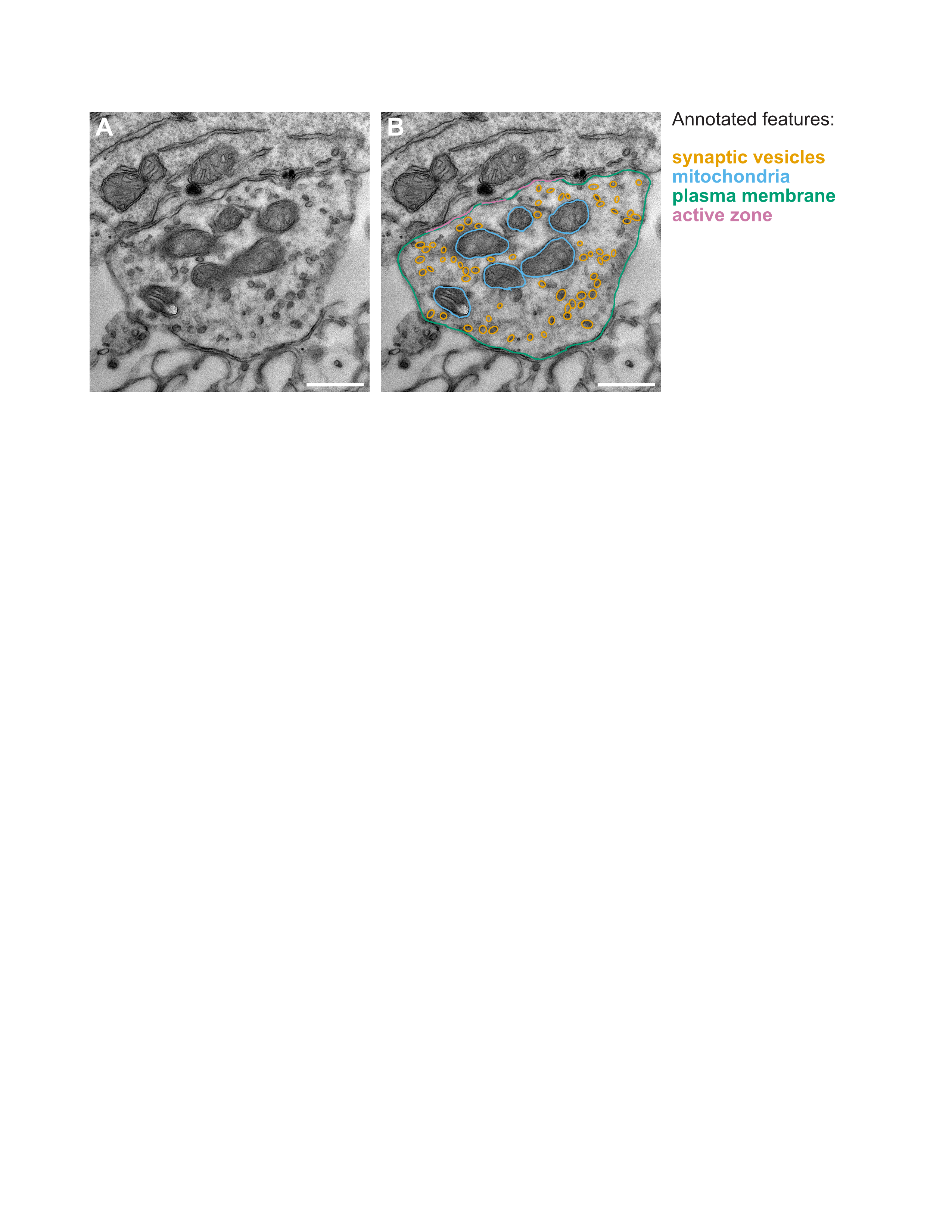

### Extended Data for Figure 3

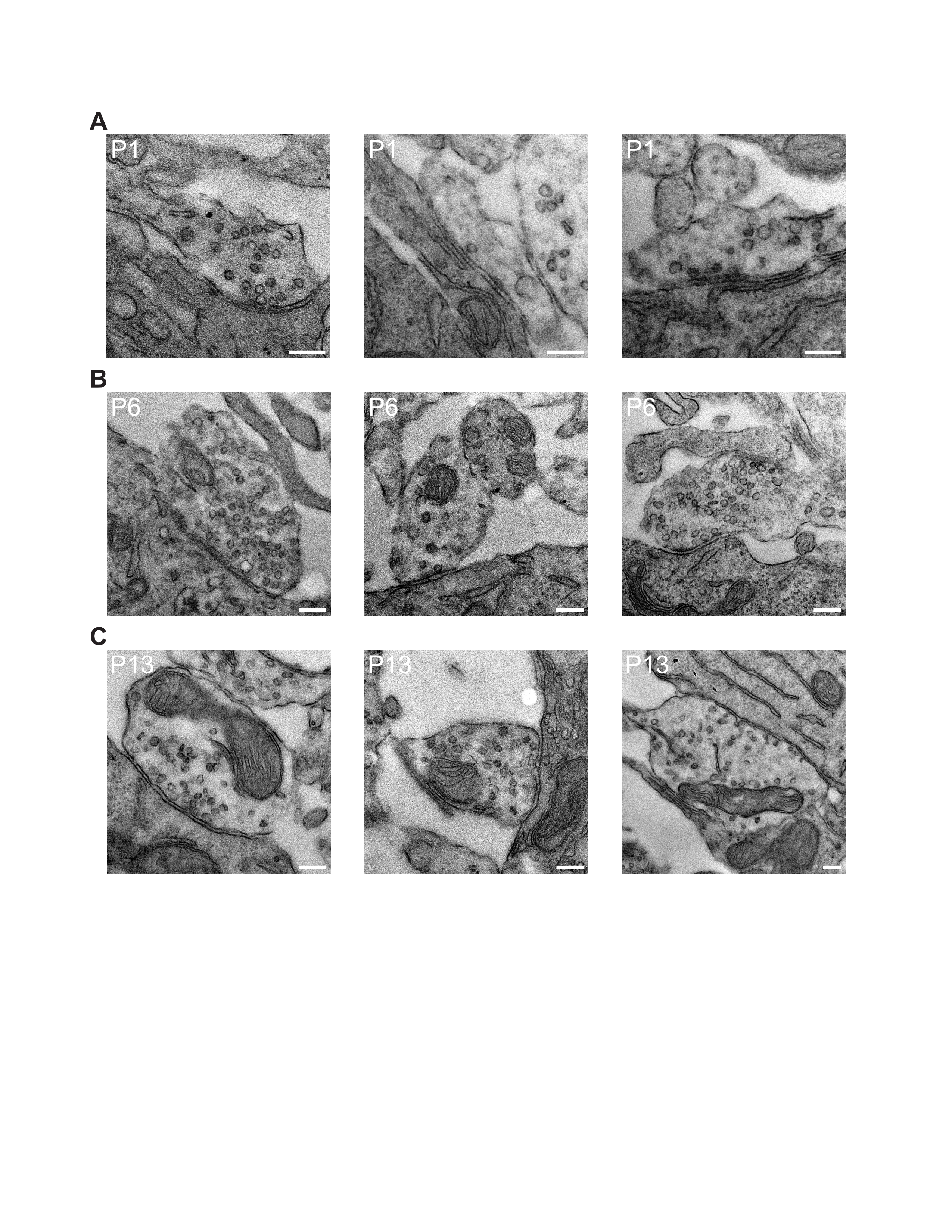

### Extended Data for Figure 6

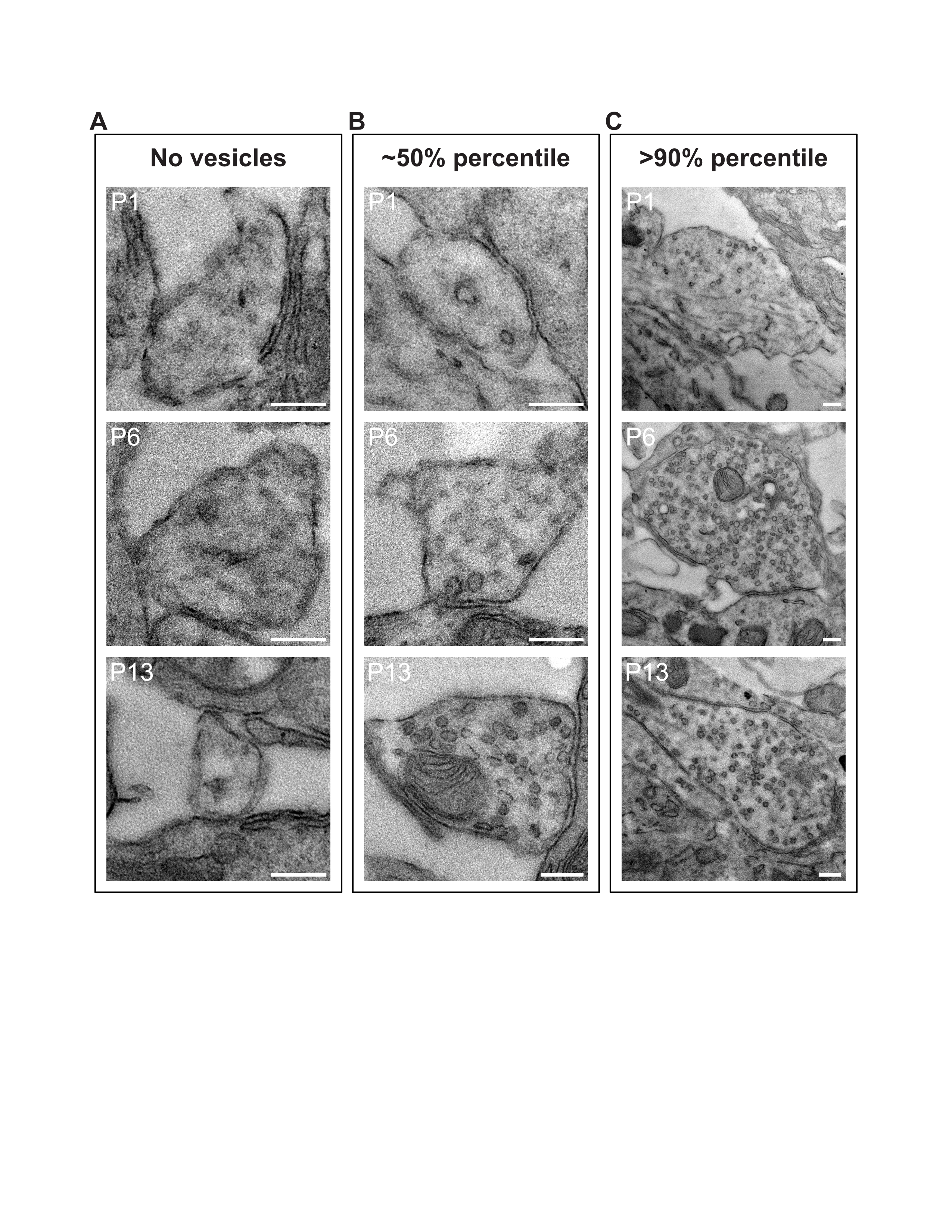

### List of antibodies and stains used for immunohistochemistry

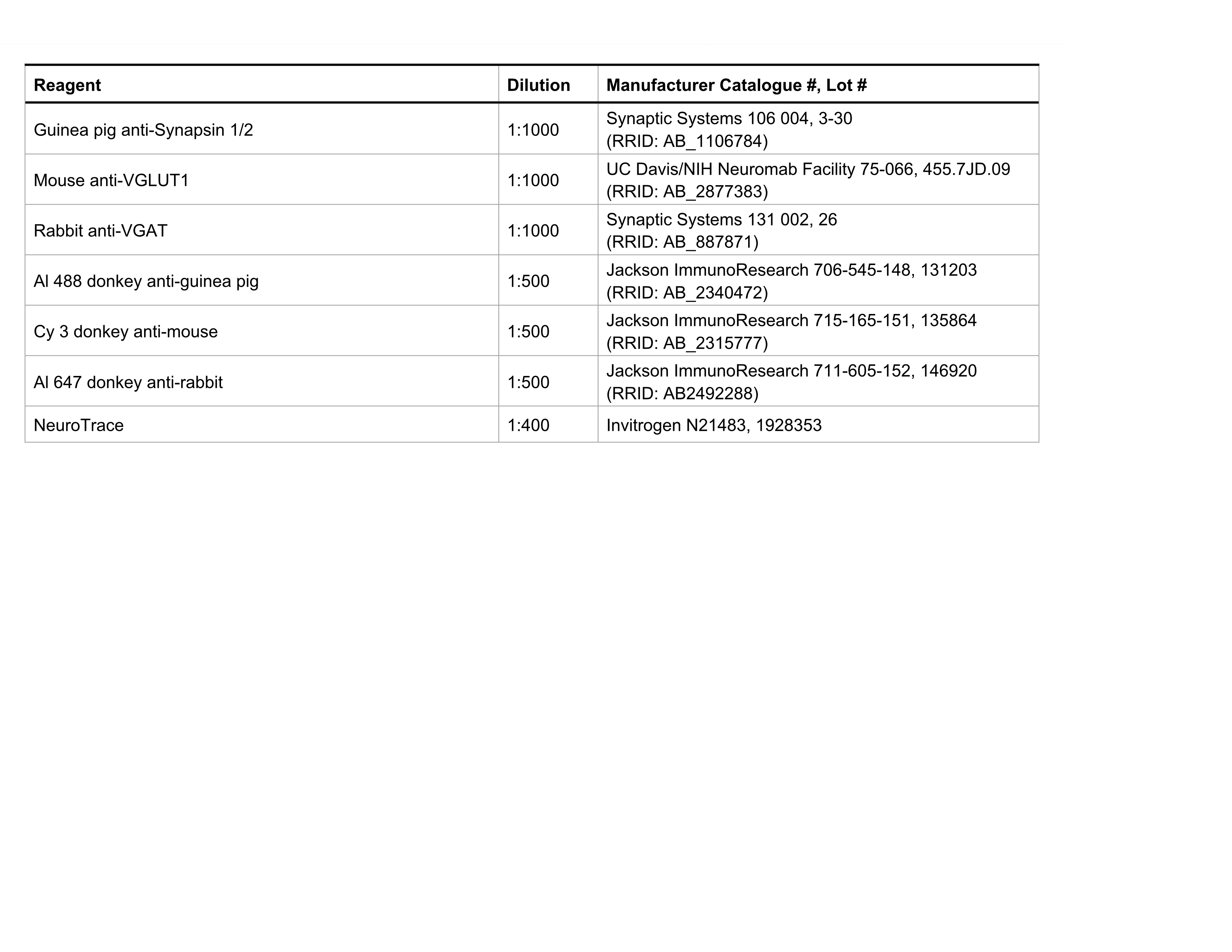
